# Multi-allelic Exact tests for Hardy-Weinberg equilibrium that account for gender

**DOI:** 10.1101/172874

**Authors:** Jan Graffelman, Bruce S. Weir

**Author notes:** Corresponding author: Jan Graffelman, Department of Statistics and Operations Research, Universitat Politècnica de Catalunya, Avinguda Diagonal 647, 08028 Barcelona, Spain.

## Abstract

Statistical tests for Hardy-Weinberg equilibrium are important elementary tools in genetic data analysis. X-chromosomal variants have long been tested by applying autosomal test procedures to females only, and gender is usually not considered when testing autosomal variants for equilibrium. Recently, we proposed specific X-chromosomal exact test procedures for bi-allelic variants that include the hemizygous males, as well as autosomal tests that consider gender. In this paper we present the extension of the previous work for variants with multiple alleles. A full enumeration algorithm is used for the exact calculations of tri-allelic variants. For variants with many alternate alleles we use a permutation test. Some empirical examples with data from the 1000 genomes project are discussed.

## 1 Introduction

Testing genetic variants for Hardy-Weinberg proportions (HWP) is an important part of many genetic studies. Genetic markers are, in general, in the absence of disturbing forces, expected to have genotype frequencies that correspond to HWP. Hardy-Weinberg proportions are often assumed in, among others, basic models in genetic epidemiology (E.g. the alleles test (Laird and Lange, 2011)), in relatedness estimation by maximum likelihood (Thompson, 1975), and in calculations in forensic genetics (Evett and Weir, 1998). In modern association studies, genetic variants are tested for equilibrium with exact procedures on a genome-wide scale, mainly for quality control purposes with the aim of identifying variants susceptible to genotyping errors. Inference on HWP for the X-chromosome is complicated by the fact that males have only one copy. Until recently, Hardy-Weinberg equilibrium on the X-chromosome has therefore been tested by using females only. In recent work (Graffelman and Weir, 2016) we proposed a modification of the exact test for Hardy-Weinberg equilibrium for bi-allelic X chromosomal variants, and designed an exact procedure that simultaneously tests for Hardy-Weinberg proportions in females and equality of allele frequencies (EAF) in the sexes. In subsequent work, we also proposed statistical procedures that account for gender when testing for HWP at bi-allelic autosomal variants (Graffelman and Weir, 2017). The number of polymorphisms used in modern genetic studies has increased tremendously over the years, and consequently more multi-allelic variants such as indels and micro-satellites, have been discovered. HWP tests for multi-allelic variants have been studied by several authors. The classical result stems from Levene (1949), who proposed an exact test for HWP of multi-allelic variants. Chapco (1976) considered an alternative exact test for the bi and tri-allelic case, based on the idea of Edwards and Cannings (1969) of distinguishing male and female gametes. A computer-intensive complete enumeration algorithm for Levene’s multi-allelic exact test was given by Louis and Dempster (1987). Computationally more efficient algorithms for determining the p-value of the multi-allelic exact tests have been developed by Guo and Thompson (1992) who used both a permutation and a Markov-Chain approach. Huber et al. (2006) proposed a method for faster generation of permuted data sets. Engels (2009) achieved speed improvements in exact calculations by using network algorithms. However, multi-allelic exact test procedures that account for gender, both for the X-chromosome, and for the autosomes, are currently not available. In this contribution we give the extension of our previous results (Graffelman and Weir, 2016, 2017) for the case of multiple alleles. We propose an exact procedure that is a straightforward extension of the probability density given by Levene (1949). The structure of the paper is as follows. In Section 2 we derive the multi-allelic exact tests that account for gender. In Section 3 we discuss a small artificial example to illustrate the calculations with complete enumeration for the tri-allelic case. We use permutation methods for estimating exact p-values for variants with larger numbers of alleles. Section 4 gives some empirical examples with tri and multi-allelic indels and SNVs taken from the 1000 Genomes project (The 1000 Genomes Project Consortium, 2015). A discussion (Section 5) finishes the paper.

## 2 Theory

We briefly review multi-allelic autosomal exact inference as developed by Levene (1949), and proceed to derive the probability densities that account for gender for both autosomal and X-chromosomal variants. We consider a variant with *k* alleles *a*_1_,*a*_2_, …*a_k_*, and let *n_i_* represent the total sample count of the *i*th allele, with *i* = 1,…, *k*. If the sexes are not distinguished, we use *n_ij_* with *i* ≥ *j* to refer to the total number of *a_i_a_j_* genotypes, including males and females. Thus, *n_ii_* refers to a homozygote *a_i_a_j_*, whereas *n_ij_* with *i* > *j* refers to a heterozygote *a_i_a_j_*. We generally represent the data in a lower triangular matrix as shown in Table 1.

**Table 1:**
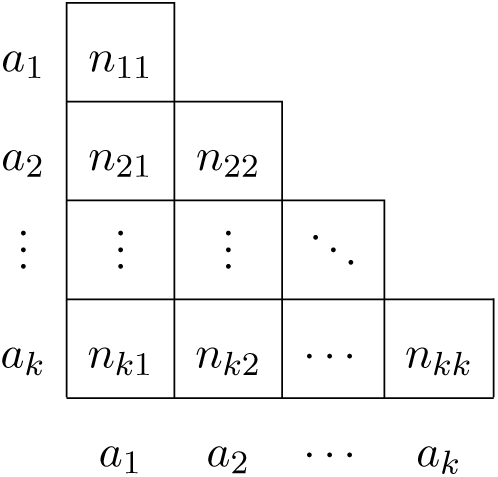
Lower triangular matrix layout for autosomal genotype counts.

|  |  |  |  |  |
| --- | --- | --- | --- | --- |
| $a_1$ | $n_{11}$ | | | |
| $a_2$ | $n_{21}$ | $n_{22}$ | | |
| ⋮ | ⋮ | ⋮ | $\ddots$ | |
| $a_k$ | $n_{k1}$ | $n_{k2}$ | ⋯ | $n_{kk}$ |
| | $a_1$ | $a_2$ | ⋯ | $a_k$ |

We let *n* represent the sample size (number of individuals) and *n_t_* the total number of alleles, with *n_t_* = 2*n* in the autosomal case.

For X-chromosomal exact procedures that distinguish gender, we let *n_mi_* be the observed number of hemizygous males with genotype *a_i_*, such that the number of males *n_m_* is given by
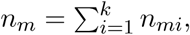
and we use *n_fij_* to represent the female genotype counts. The total sample size is given by *n* = *n_m_* + *n_f_*, and the total number of alleles is given by *n_t_* = *n_m_* + 2*n_f_*. The data are now represented by a vector for males, and a triangular matrix for females, as shown in Table 2.

**Table 2:** Layout of X-chromosomal genotype counts for males (vector) and females (lower triangular matrix).

|  |  |  |  |  |  |  |
| --- | --- | --- | --- | --- | --- | --- |
| $a_1$ | $n_{m1}$ | $a_1$ | $n_{f11}$ | | | |
| $a_2$ | $n_{m2}$ | $a_2$ | $n_{f21}$ | $n_{f22}$ | | |
| ⋮ | ⋮ | ⋮ | ⋮ | ⋮ | $\ddots$ | |
| $a_k$ | $n_{mk}$ | $a_k$ | $n_{fk1}$ | $n_{fk2}$ | ⋯ | $n_{fkk}$ |
| | | | $a_1$ | $a_2$ | ⋯ | $a_k$ |

Finally, for autosomal procedures that take gender into account, we let *n_fij_* be the observed number of female *a_i_a_j_* genotypes, with
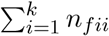
being the total female homozygote count and ∑_*i*>*j*_ *n_fij_* the total female heterozygote count. The number of females *n_f_* is then given by
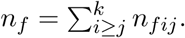
The obvious analogous quantities for males are *n_mij_* and *n_m_*. The total number of alleles is given by *n_t_* = 2(*n_m_* + *n_f_*), and the data are now represented by two triangular matrices as shown in Table 3.

**Table 3:** Two-fold triangular matrix layout of autosomal genotype counts accounting for gender, with male and female genotype counts.

|  |  |  |  |  |
| --- | --- | --- | --- | --- |
| $a_1$ | $n_{m11}$ | | | |
| $a_2$ | $n_{m21}$ | $n_{m22}$ | | |
| ⋮ | ⋮ | ⋮ | $\ddots$ | |
| $a_k$ | $n_{mk1}$ | $n_{mk2}$ | ⋯ | $n_{mkk}$ |
| | $a_1$ | $a_2$ | ⋯ | $a_k$ |
| $a_1$ | $n_{f11}$ | | | |
| $a_2$ | $n_{f21}$ | $n_{f22}$ | | |
| ⋮ | ⋮ | ⋮ | $\ddots$ | |
| $a_k$ | $n_{fk1}$ | $n_{fk2}$ | ⋯ | $n_{fkk}$ |
| | $a_1$ | $a_2$ | ⋯ | $a_k$ |

### 2.1 Classical autosomal exact inference

Exact inference for autosomal variants with multiple alleles is based on the conditional distribution of the genotype counts, considering all observed allele counts as given. This distribution was derived by Levene (1949), and is given by

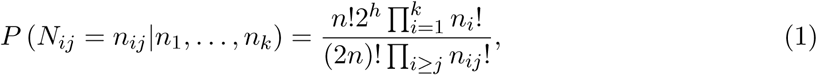

where *h* = ∑_*i*>*j*_ *n_ij_* is the total heterozygote frequency. We re-derive Levene’s density, but taking gender into account, for the X chromosome in Section 2.2, and for the autosomes in Section 2.3.

### 2.2 X chromosomal exact inference with gender

We condition on the numbers of males (*n_m_*) and females (*n_f_*). Under the hypothesis of Hardy-Weinberg equilibrium in females and equality of allele frequencies in the sexes, the distribution of the genotype counts is given by

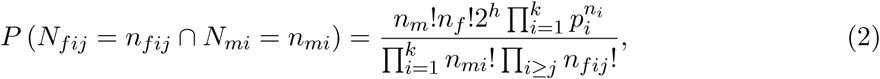

where *h* = ∑_*i*>*j*_*n_fij_* is the total female heterozygote count, and *p_i_* the relative frequency of the *i*th allele. Under HWP and EAF, the allele counts *N_i_* follow a multinomial distribution given by

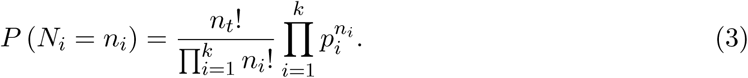

The conditional distribution of the genotype counts given the allele counts is obtained by dividing Equation (2) by (3), and is given by:

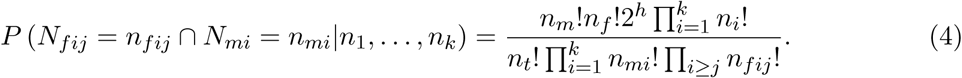

We note that if the male allele and genotype counts are set to zero, then (4) reduces to Levene’s density (1), but applied to the females only. We also note that if counts involving alleles beyond the second one (*j* > 2) are set to zero, the Graffelman-Weir density (2016) for the bi-allelic case is obtained.

### 2.3 Autosomal exact inference with gender

We again condition on the number of males (*n_m_*) and females (*n_f_*). Under the hypothesis of Hardy-Weinberg equilibrium and equality of allele frequencies in the sexes, the distribution of the genotype counts is given by

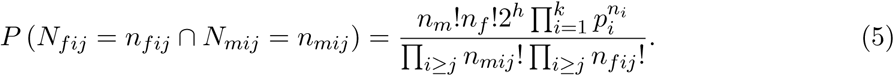

where *h* is total number of heterozygotes *h* = ∑_*i*>*j*_ (*n_mij_* + *n_fij_*). The distribution of the allele counts is again given by the multinomial distribution. Dividing (5) by (3) we obtain the conditional density

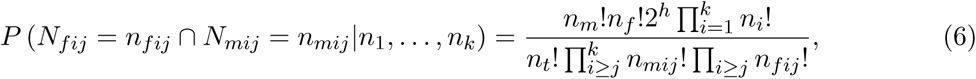

which can be used for exact inference for HWP while accounting for gender. In the remainder, we will generally refer to the exact tests based on Equations (4) and (6), that have a composite null hypothesis, HWP *and* EAF, as omnibus exact tests.

## 3 Artificial data example

In this Section we discuss an artificial data example to illustrate the calculations of an X-chromosomal multi-allelic exact test that takes sex into account. In order to calculate the p-value of the test, we use an algorithm that enumerates all possible outcomes. For this purpose we extend the algorithm described by Louis and Dempster (1987). The algorithm has input parameters *n_A_*, *n_B_*, *n_C_*, *n_m_* and *n_f_*, and *n_f_*, and assumes *n_A_* ≤ *n_B_* ≤ *n_C_*, and it proceeds by first assigning the maximum possible amount of minor A alleles to males, and then assigns the remaining alleles to the females. All possible outcomes for the female genotype counts are generated by the Louis-Dempster algorithm. We note that the overall minor allele does not necessarily coincide with the minor allele in females. For this reason, alleles allocated to females are first sorted and later re-labeled to ensure consistency with the original labeling of the alleles. The algorithm is readily extended for additional alleles. Each additional allele implies two extra for loops, one for the male genotypes and another one for the female genotypes. Table 4 shows all possible outcomes for a sample with genotype counts (*A* = 2, *B* = 2, *C* = 2, *AA* = 0, *AB* = 1, *AC* = 0, *BB* = 0, *BC* = 2, *CC* = 1), consisting of 10 individuals (6 males and 4 females) with allele counts *A* = 3, *B* = 5 and *C* = 6. The samples are ordered as they are produced by the algorithm. Table 4 shows how initially all three minor A alleles are first assigned to A males, leaving 6-3 = 3 second minor B alleles for B males and 0 C alleles for C males. This leaves 0 A, 2 B and 6 C alleles to be assigned to females. All outcomes for females with these allele counts are generated by the Louis-Dempster algorithm. Next, the number of minor B alleles in males is decreased by one, and the number of C males increased by 1. This continues till all possible male genotype counts are exhausted. For each possible set of male genotype counts, the remaining alleles are assigned to females, in all possible ways, so creating repeated entries of the male genotype counts in the table. Note that even for a small sample of 10 individuals, 75 outcomes are possible. If sex would have been ignored, then there would be only 1 A, 3 B and 4 C to be assigned to females, and only 4 outcomes are possible.

**Table 4:** Table of all possible outcomes and probabilities of a sample consisting of 10 individuals (6 males and 4 females) with total allele counts *A* = 3, *B* = 5 and *C* = 6 (first part).

| Nr. | Males |  |  | Females |  |  |  |  |  | Prob. |
| --- | --- | --- | --- | --- | --- | --- | --- | --- | --- | --- |
|  | A | B | C | AA | AB | AC | BB | BC | CC |  |
| 1 | 3 | 3 | 0 | 0 | 0 | 0 | 0 | 2 | 2 | 0.0029 |
| 2 | 3 | 3 | 0 | 0 | 0 | 0 | 1 | 0 | 3 | 0.0005 |
| 3 | 3 | 2 | 1 | 0 | 0 | 0 | 0 | 3 | 1 | 0.0114 |
| 4 | 3 | 2 | 1 | 0 | 0 | 0 | 1 | 1 | 2 | 0.0086 |
| 5 | 3 | 1 | 2 | 0 | 0 | 0 | 0 | 4 | 0 | 0.0057 |
| 6 | 3 | 1 | 2 | 0 | 0 | 0 | 1 | 2 | 1 | 0.0171 |
| 7 | 3 | 1 | 2 | 0 | 0 | 0 | 2 | 0 | 2 | 0.0021 |
| 8 | 3 | 0 | 3 | 0 | 0 | 0 | 1 | 3 | 0 | 0.0038 |
| 9 | 3 | 0 | 3 | 0 | 0 | 0 | 2 | 1 | 1 | 0.0029 |
| 10 | 2 | 4 | 0 | 0 | 1 | 0 | 0 | 0 | 3 | 0.0007 |
| 11 | 2 | 4 | 0 | 0 | 0 | 1 | 0 | 1 | 2 | 0.0043 |
| 12 | 2 | 3 | 1 | 0 | 1 | 0 | 0 | 1 | 2 | 0.0171 |
| 13 | 2 | 3 | 1 | 0 | 0 | 1 | 0 | 2 | 1 | 0.0343 |
| 14 | 2 | 3 | 1 | 0 | 0 | 1 | 1 | 0 | 2 | 0.0086 |
| **15** | **2** | **2** | **2** | **0** | **1** | **0** | **0** | **2** | **1** | **0.0514** |
| 16 | 2 | 2 | 2 | 0 | 1 | 0 | 1 | 0 | 2 | 0.0128 |
| 17 | 2 | 2 | 2 | 0 | 0 | 1 | 0 | 3 | 0 | 0.0343 |
| 18 | 2 | 2 | 2 | 0 | 0 | 1 | 1 | 1 | 1 | 0.0514 |
| 19 | 2 | 1 | 3 | 0 | 0 | 1 | 1 | 2 | 0 | 0.0343 |
| 20 | 2 | 1 | 3 | 0 | 0 | 1 | 2 | 0 | 1 | 0.0086 |
| 21 | 2 | 1 | 3 | 0 | 1 | 0 | 0 | 3 | 0 | 0.0228 |
| 22 | 2 | 1 | 3 | 0 | 1 | 0 | 1 | 1 | 1 | 0.0343 |
| 23 | 2 | 0 | 4 | 0 | 0 | 1 | 2 | 1 | 0 | 0.0043 |
| 24 | 2 | 0 | 4 | 0 | 1 | 0 | 1 | 2 | 0 | 0.0086 |
| 25 | 2 | 0 | 4 | 0 | 1 | 0 | 2 | 0 | 1 | 0.0021 |
| 26 | 1 | 5 | 0 | 0 | 0 | 2 | 0 | 0 | 2 | 0.0009 |
| 27 | 1 | 5 | 0 | 1 | 0 | 0 | 0 | 0 | 3 | 0.0001 |
| 28 | 1 | 4 | 1 | 0 | 1 | 1 | 0 | 0 | 2 | 0.0086 |
| 29 | 1 | 4 | 1 | 0 | 0 | 2 | 0 | 1 | 1 | 0.0171 |
| 30 | 1 | 4 | 1 | 1 | 0 | 0 | 0 | 1 | 2 | 0.0043 |
| 31 | 1 | 3 | 2 | 0 | 2 | 0 | 0 | 0 | 2 | 0.0086 |
| 32 | 1 | 3 | 2 | 0 | 1 | 1 | 0 | 1 | 1 | 0.0685 |
| 33 | 1 | 3 | 2 | 0 | 0 | 2 | 0 | 2 | 0 | 0.0343 |
| 34 | 1 | 3 | 2 | 0 | 0 | 2 | 1 | 0 | 1 | 0.0171 |
| 35 | 1 | 3 | 2 | 1 | 0 | 0 | 0 | 2 | 1 | 0.0171 |
| 36 | 1 | 3 | 2 | 1 | 0 | 0 | 1 | 0 | 2 | 0.0043 |
| 37 | 1 | 2 | 3 | 0 | 2 | 0 | 0 | 1 | 1 | 0.0343 |
| 38 | 1 | 2 | 3 | 0 | 1 | 1 | 0 | 2 | 0 | 0.0685 |

**Table 5:** Table of all possible outcomes and probabilities of a sample consisting of 10 individuals (6 males and 4 females) with total allele counts *A* = 3, *B* = 5 and *C* = 6 (second part).

| Nr. | Males |  |  | Females |  |  |  |  |  | Prob. |
| --- | --- | --- | --- | --- | --- | --- | --- | --- | --- | --- |
|  | A | B | C | AA | AB | AC | BB | BC | CC |  |
| 39 | 1 | 2 | 3 | 0 | 1 | 1 | 1 | 0 | 1 | 0.0343 |
| 40 | 1 | 2 | 3 | 0 | 0 | 2 | 1 | 1 | 0 | 0.0343 |
| 41 | 1 | 2 | 3 | 1 | 0 | 0 | 0 | 3 | 0 | 0.0114 |
| 42 | 1 | 2 | 3 | 1 | 0 | 0 | 1 | 1 | 1 | 0.0171 |
| 43 | 1 | 1 | 4 | 0 | 0 | 2 | 2 | 0 | 0 | 0.0043 |
| 44 | 1 | 1 | 4 | 0 | 1 | 1 | 1 | 1 | 0 | 0.0343 |
| 45 | 1 | 1 | 4 | 0 | 2 | 0 | 0 | 2 | 0 | 0.0171 |
| 46 | 1 | 1 | 4 | 0 | 2 | 0 | 1 | 0 | 1 | 0.0086 |
| 47 | 1 | 1 | 4 | 1 | 0 | 0 | 1 | 2 | 0 | 0.0086 |
| 48 | 1 | 1 | 4 | 1 | 0 | 0 | 2 | 0 | 1 | 0.0021 |
| 49 | 1 | 0 | 5 | 0 | 1 | 1 | 2 | 0 | 0 | 0.0017 |
| 50 | 1 | 0 | 5 | 0 | 2 | 0 | 1 | 1 | 0 | 0.0034 |
| 51 | 1 | 0 | 5 | 1 | 0 | 0 | 2 | 1 | 0 | 0.0009 |
| 52 | 0 | 5 | 1 | 0 | 0 | 3 | 0 | 0 | 1 | 0.0011 |
| 53 | 0 | 5 | 1 | 1 | 0 | 1 | 0 | 0 | 2 | 0.0009 |
| 54 | 0 | 4 | 2 | 0 | 1 | 2 | 0 | 0 | 1 | 0.0086 |
| 55 | 0 | 4 | 2 | 1 | 1 | 0 | 0 | 0 | 2 | 0.0021 |
| 56 | 0 | 4 | 2 | 0 | 0 | 3 | 0 | 1 | 0 | 0.0057 |
| 57 | 0 | 4 | 2 | 1 | 0 | 1 | 0 | 1 | 1 | 0.0086 |
| 58 | 0 | 3 | 3 | 0 | 2 | 1 | 0 | 0 | 1 | 0.0114 |
| 59 | 0 | 3 | 3 | 0 | 1 | 2 | 0 | 1 | 0 | 0.0228 |
| 60 | 0 | 3 | 3 | 1 | 1 | 0 | 0 | 1 | 1 | 0.0114 |
| 61 | 0 | 3 | 3 | 1 | 0 | 1 | 0 | 2 | 0 | 0.0114 |
| 62 | 0 | 3 | 3 | 0 | 0 | 3 | 1 | 0 | 0 | 0.0038 |
| 63 | 0 | 3 | 3 | 1 | 0 | 1 | 1 | 0 | 1 | 0.0057 |
| 64 | 0 | 2 | 4 | 0 | 1 | 2 | 1 | 0 | 0 | 0.0086 |
| 65 | 0 | 2 | 4 | 0 | 2 | 1 | 0 | 1 | 0 | 0.0171 |
| 66 | 0 | 2 | 4 | 1 | 0 | 1 | 1 | 1 | 0 | 0.0086 |
| 67 | 0 | 2 | 4 | 1 | 1 | 0 | 0 | 2 | 0 | 0.0086 |
| 68 | 0 | 2 | 4 | 0 | 3 | 0 | 0 | 0 | 1 | 0.0029 |
| 69 | 0 | 2 | 4 | 1 | 1 | 0 | 1 | 0 | 1 | 0.0043 |
| 70 | 0 | 1 | 5 | 0 | 2 | 1 | 1 | 0 | 0 | 0.0034 |
| 71 | 0 | 1 | 5 | 1 | 0 | 1 | 2 | 0 | 0 | 0.0009 |
| 72 | 0 | 1 | 5 | 0 | 3 | 0 | 0 | 1 | 0 | 0.0023 |
| 73 | 0 | 1 | 5 | 1 | 1 | 0 | 1 | 1 | 0 | 0.0034 |
| 74 | 0 | 0 | 6 | 0 | 3 | 0 | 1 | 0 | 0 | 0.0002 |
| 75 | 0 | 0 | 6 | 1 | 1 | 0 | 2 | 0 | 0 | 0.0001 |

A similar enumeration algorithm can be used for an autosomal exact test that accounts for gender. In the tri-allelic case, one can first generate all possible outcomes for the male A, B and C allele counts, given the observed number of A, B and C alleles, and the observed number of males. All possible male genotype counts are then obtained by applying the Louis-Dempster algorithm to each set of male (A,B,C) counts. A table of all possible female allele counts is obtained by subtracting the male table from the total observed allele counts. All possible female genotype counts are obtained by applying the Louis-Dempster algorithm to the female allele counts. Finally, the table of all possible genotype outcomes for the 12 genotypes can be formed by taking the Cartesian product of the male and female genotype table.

As the example shows, many tied outcomes that have the same probability arise. The observed sample (row 15) has probability 0.05138. The sum of the probabilities of all samples equally or less likely gives the p-value of the test: 0.86299. Graffelman and Moreno (2013) have advocated the mid-p value (half the probability of the observed data plus the sum of the probabilities of all extremer samples) which for this example is 0.83731, and HWP can not be rejected for this example. If, as has been the standard practice, HWP are tested by an exact test of the females only, then the test is not significant either (p = 1.00000). We note that sample 18 is a tied outcome whose probability is included in the sum that makes up the p-value.

## 4 Empirical data examples

We use data from the Japanese (JPT) sample of the 1000 Genomes project to illustrate our results. This sample consists of 56 males and 48 females. Data stored in variant call format (VCF) were downloaded from the 1000 Genomes project (http://www.internationalgenome.org/). Statistical analysis was carried out in the R environment (R Core Team, 2014). VCF files were processed in R with the VCFR package (Knaus and Grünwald, 2017). We analyze multi-allelic X chromosomal variants in Section 4.1 and autosomal variants from chromosome 7 in Section 4.2.

### 4.1 X chromosomal variants

We extracted multi-allelic variants of the X chromosome of the JPT sample. Multi-allelic variants on X are rare. Of all 3.5M variants, 87.57% were monomorphic, 12.32% were bi-allelic, 0.10% were tri-allelic, and 0.01% had four or more alleles. We consider some examples of triallelic X-chromosomal variants. Table 6 shows genotype counts and p-values of statistical tests for five tri-allelic variants. These variants are registered as indels with at least two alternate alleles.

**Table 6:** Identifier, position (in bp), genotype counts and exact p-values for five tri-allelic X chromosomal variants. HWP & EAF: omnibus exact test for HWP and EAF jointly, Perm.: approximation of the omnibus p-value by a permutation procedure with 20.000 draws, HWP (F): exact test for HWP in females only, EAF: Fisher exact test for equality of allele frequencies in males and females.

| Variant | Position | Genotype counts |  |  |  |  |  |  |  |  |
| --- | --- | --- | --- | --- | --- | --- | --- | --- | --- | --- |
|  |  | Males |  |  | Females |  |  |  |  |  |
|  |  | A | B | C | AA | AB | AC | BB | BC | CC |
| NA | 18892613 | 0 | 0 | 56 | 0 | 0 | 2 | 0 | 14 | 32 |
| NA | 44317003 | 2 | 2 | 52 | 0 | 0 | 0 | 0 | 0 | 48 |
| rs111463470 | 3107933 | 1 | 21 | 34 | 0 | 1 | 0 | 8 | 24 | 15 |
| rs79878783 | 67482671 | 0 | 15 | 41 | 0 | 0 | 1 | 3 | 41 | 3 |
| rs373113553 | 83676643 | 15 | 17 | 24 | 4 | 2 | 13 | 6 | 19 | 4 |

Table 6: Identifier, position (in bp), genotype counts and exact p-values for five tri-allelic X chromosomal variants.
| Variant | Position | Exact p-values |  |  |  |
| --- | --- | --- | --- | --- | --- |
|  |  | HWP & EAF | Perm. | HWP (F). | EAF |
| NA | 18892613 | 0.0026 | 0.0029 | 0.6923 | 0.0008 |
| NA | 44317003 | 0.0422 | 0.0418 | 1.0000 | 0.0172 |
| rs111463470 | 3107933 | 0.8309 | 0.8327 | 0.6443 | 0.7623 |
| rs79878783 | 67482671 | 0.0000 | 0.0000 | 0.0000 | 0.0079 |
| rs373113553 | 83676643 | 0.0310 | 0.0304 | 0.0072 | 0.8880 |

We tested these variants for HWP by using a multi-allelic exact test on the females only (HWP (F)), and by the multi-allelic exact procedure developed in this paper, using both males and females. The joint exact test (HWP $ EAF) was also carried out by avoiding the complete enumeration and using 20.000 permutations to estimate the exact p-value. The obtained permutation test p-values are seen to be close to the exact p-values. We also tested equality of the three allele frequencies (EAF) in the sexes by using a Fisher exact test on the 2 × 3 cross table of sex by allele counts.

Variant at position 18892613 (without identifier) is monomorphic in males, but it has all three alleles in females, as both major-allele involving heterozygotes AC and BC females are found. An exact test using females only does not reject HWP. The joint exact test does reject the joint null of HWP & EAF. EAF is also rejected marginally by Fisher’s exact test. This variant has an unexpected pattern of genotype counts that goes undetected if HWP are tested in females alone.

Similarly, a variant at position 44317003 (without identifier) is monomorphic in females, but all three alleles are observed in males. An exact test for HWP in females has p-value 1, but the joint test rejects the joint null of HWP & EAF. A Fisher exact test for EAF is significant. Again, the variant has an unexpected pattern of genotype counts that goes undetected if HWP are tested in females alone.

Variant rs111463470 has low minor (A) allele frequency in both males and females, B allele frequencies are intermediate and C allele frequencies are the largest in both sexes. All exact tests (HWP & EAF, HWP (F), EAF) are clearly non-significant. This variant represents a pattern of genotype counts that is common in the data.

Variant rs79878783 significant in all exact tests. The pattern of the variant is unexpected in the sense that the B allele is more frequent in females than in males, and that there is an excess of BC heterozygotes. For this case, the male allele frequencies suggest that part of the female BC heterozygotes may in fact be CC homozygotes.

Variant rs373113553 is highly polymorphic. There is no significant difference in allele frequencies between the sexes. Disequilibrium is due to a lack of AB heterozygotes.

We proceed to analyze all 2979 tri-allelic X-chromosomal variants found, and represent them using four plots represented in Figure 1.

**Figure 1:**
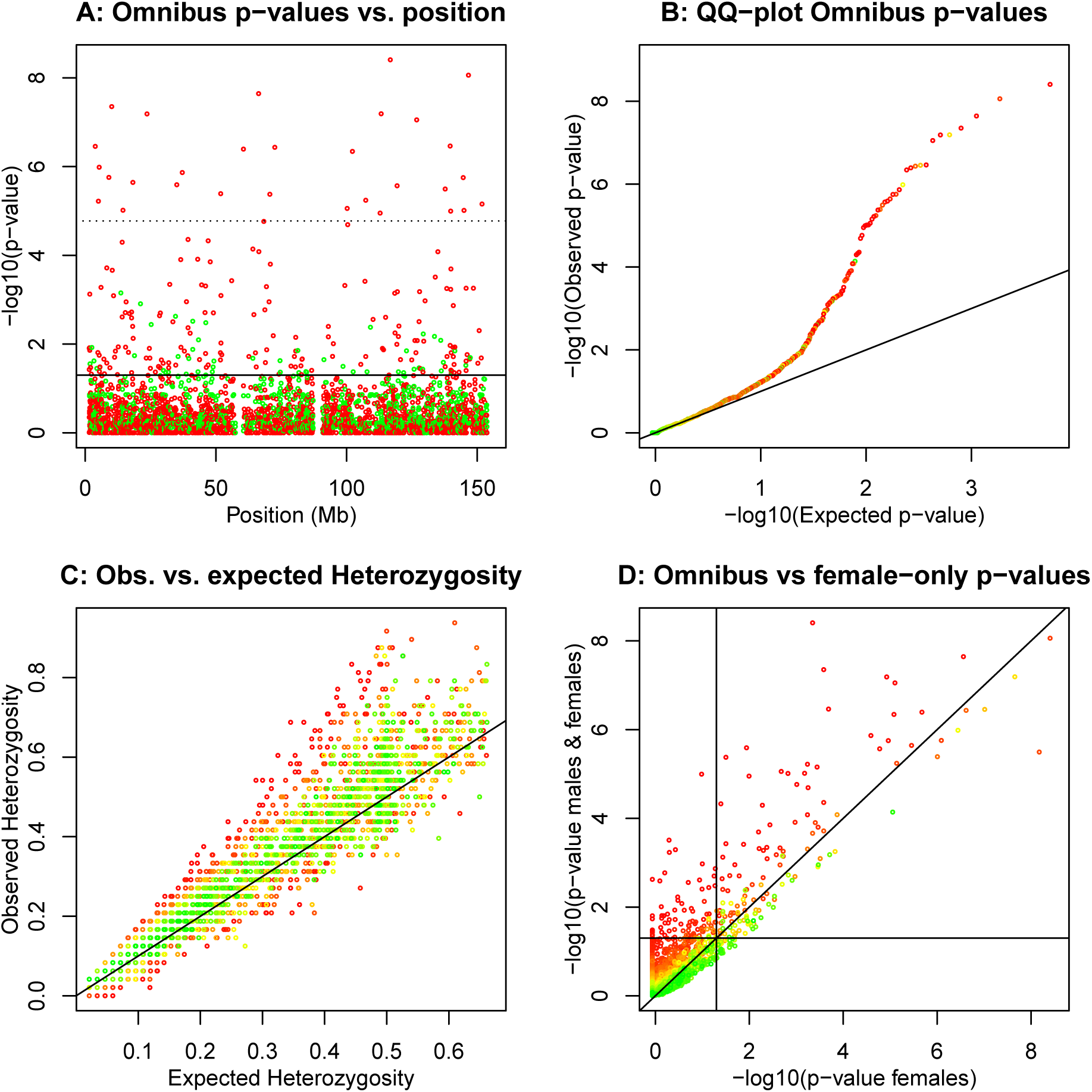
Hardy-Weinberg equilibrium tests for 2979 tri-allelic X chromosomal variants. A: p-values of the omnibus exact test (HWP & EAF) versus variant position. The solid horizontal line corresponds to *α* = 0.05, and the dotted horizontal line corresponds to the Bonferroni corrected threshold. Variants colour-coded according to having excess of heterozygotes (red) or lack of heterozygotes (green) with respect to HWP. B: QQ-plot of the omnibus exact p-values. C: Observed versus expected heterozygosity. D: Omnibus exact p-values versus exact p-values obtained in a test using the females only. Horizontal and vertical lines correspond to *α* = 0.05. Panels B, C and D show variants colour-coded according to the p-value of an exact test for EAF.

Figure 1A shows that the tri-allelics are scattered along the whole X chromosome, and that the most significant variants typically have an excess of heterozygotes. Figure 1B shows a QQ-plot of the omnibus exact p-values in the logarithmic scale. There is deviation from the uniform distribution in the lower tail of the p-value distribution, and the corresponding variants typically have significantly different allele frequencies in the sexes. Figure 1C shows the observed against the expected heterozygosity. The maximum possible expected heterozygosity is 2/3 for a triallelic variant. Observed heterozygosity is often larger than the expected heterozygosity. 65% of the variants are above the line *y* = *x*, showing that the tri-allelic variants are predominantly characterized by excess heterozygosity. Variants whose observed heterozygosity is much larger or much lower than their expected heterozygosity, appear with significant colour in a test for equality of allele frequencies. The deviation from HWP in females is related to the difference in allele frequency between the sexes. Figure 1D shows a plot of omnibus exact p-values against exact p-values obtained by using females only. The omnibus test detects many variants as significant that are non-significant in a females-only test, due to differences in allele frequencies in the sexes. Likewise, variants with very similar allele frequencies in the sexes can appear significant in a females-only test, and non-significant in the omnibus. For about 4% of the variants the test result is inverted (from significant to non-significant or vice verse with *α* = 0.05). Figure 1D is similar to what has been observed for bi-allelic X chromosomal variants (Graffelman and Weir, 2016, Figure 6).

We finish the analysis of the multi-allelic variants on the X chromosome with a few examples involving more than three alleles. The genotype counts of a three, four, five and six allelic indel on chromosome X are shown in Table 7.

**Table 7:**
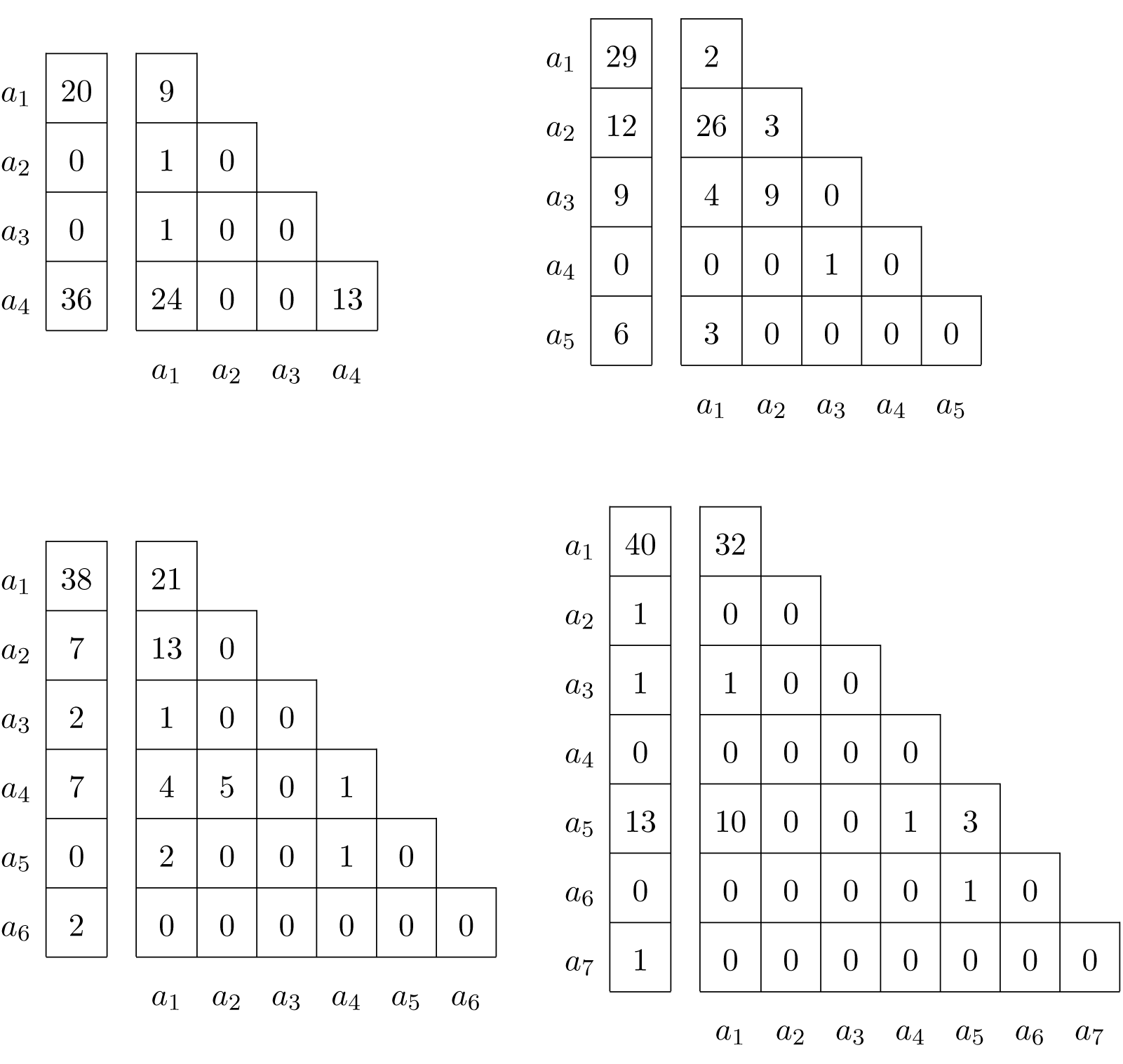
Male and female genotype counts of a four, five, six and seven allelic indel on chromosome X.

The results of several statistical tests with these multi-allelic indels are shown in Table 8. Variant rs67657605 is a four-allelic variant, with alternate allele *a*_4_ being the most common in both males and females. Reference allele *a*_1_ is second most common in both sexes. Other alternate alleles are rare. The probability of the observed sample is 0.012, and the p-value of the omnibus exact test, estimated by permutation, is 0.3101, indicating the HWP and EAF can not be rejected. Consistently, neither an exact test for HWP using females only, nor a Fisher exact test for equality of allele frequencies reject the null hypothesis. The variant is in fact close to being bi-allelic with only two rare alternate alleles. If the two rare alleles are ignored, and the variant is tested as bi-allelic, similar conclusions are obtained (HWP & EAF p-value = 0.4862; HWP (F) p-value = 1.000; EAF p-value = 0.3028).

**Table 8:** P values of statistical tests for multi-allelic indels on chromosome X. HWP & EAF: joint exact test for Hardy-Weinberg proportions and equality of allele frequencies (obtained by permutation); HWP (F): exact test for Hardy-Weinberg proportions in females only; EAF: Fisher Exact test for equality of allele frequencies in the sexes.

|  | Variant | # alleles | HWP & EAF | HWP (F) | EAF |
| --- | --- | --- | --- | --- | --- |
| 1 | rs67657605 | 4 | 0.3101 | 0.6553 | 0.4578 |
| 2 | NA | 5 | 0.0001 | 0.0002 | 0.0253 |
| 3 | rs60184331 | 6 | 0.2254 | 0.1714 | 0.2386 |
| 4 | rs59130472 | 7 | 0.1070 | 0.0497 | 0.5045 |

The five-allelic variant (which has no rs identifier) is significant for all three tests, indicating both differences in allele frequencies and deviation from Hardy-Weinberg proportions. Its reference allele *a*_1_ is the most common allele in males, whereas the the main alternate allele *a*_2_ is twice as frequent in females. The variant has a very high observed heterozygosity (*H_o_* = 0.896), whereas its expected heterozygosity, *H_e_*, is 0.647. The results suggest that the variant has some kind of genotyping problem. If the the single rare *a_4_* allele is eliminated from the sample, similar conclusions are reached (HWP & EAF p value = 0.0003; HWP p-value = 0.0003; EAF p-value = 0.0197).

The six-allelic variant rs60184331 is non-significant in all tests. This variant has three rare alleles (*a*_3_,*a*_5_ and *a*_6_). If these are ignored, a tri-allelic variant remains, for which equality of allele frequencies holds, but evidence for deviation from HWP in females is found (HWP & EAF p-value = 0.1110; HWP (F) p-value = 0.0455; EAF p-value = 0.6102).

Variant rs59130472 with seven alleles shows evidence for deviation from HWP in females. This variant has four rare alleles that mostly occur in only one of the sexes (*a*_2_,*a*_3_,*a*_4_,*a*_6_ and *a*_7_). If these are ignored, the variant is converted into a bi-allelic one, for which no significant deviations are found (HWP & EAF p-value = 0.1434; HWP (F) p-value = 0.1180; EAF p-value = 0.3908).

The distributions of the probabilities of the permuted samples of the four studied multi-allelic variants are shown in Figure 2. Because many outcomes have a small probability, this distribution is more conveniently displayed in a logarithmic scale. The p-values of the permutation test for the joint HWP & EAF test correspond to areas in the right tail of this distribution, where the probability of the observed sample is indicated by a vertical line. These distributions give a graphical appraisal of how extreme the observed sample is under the joint assumption of HWP and EAF, and clearly illustrate the significance of the five-allelic indel.

**Figure 2:**
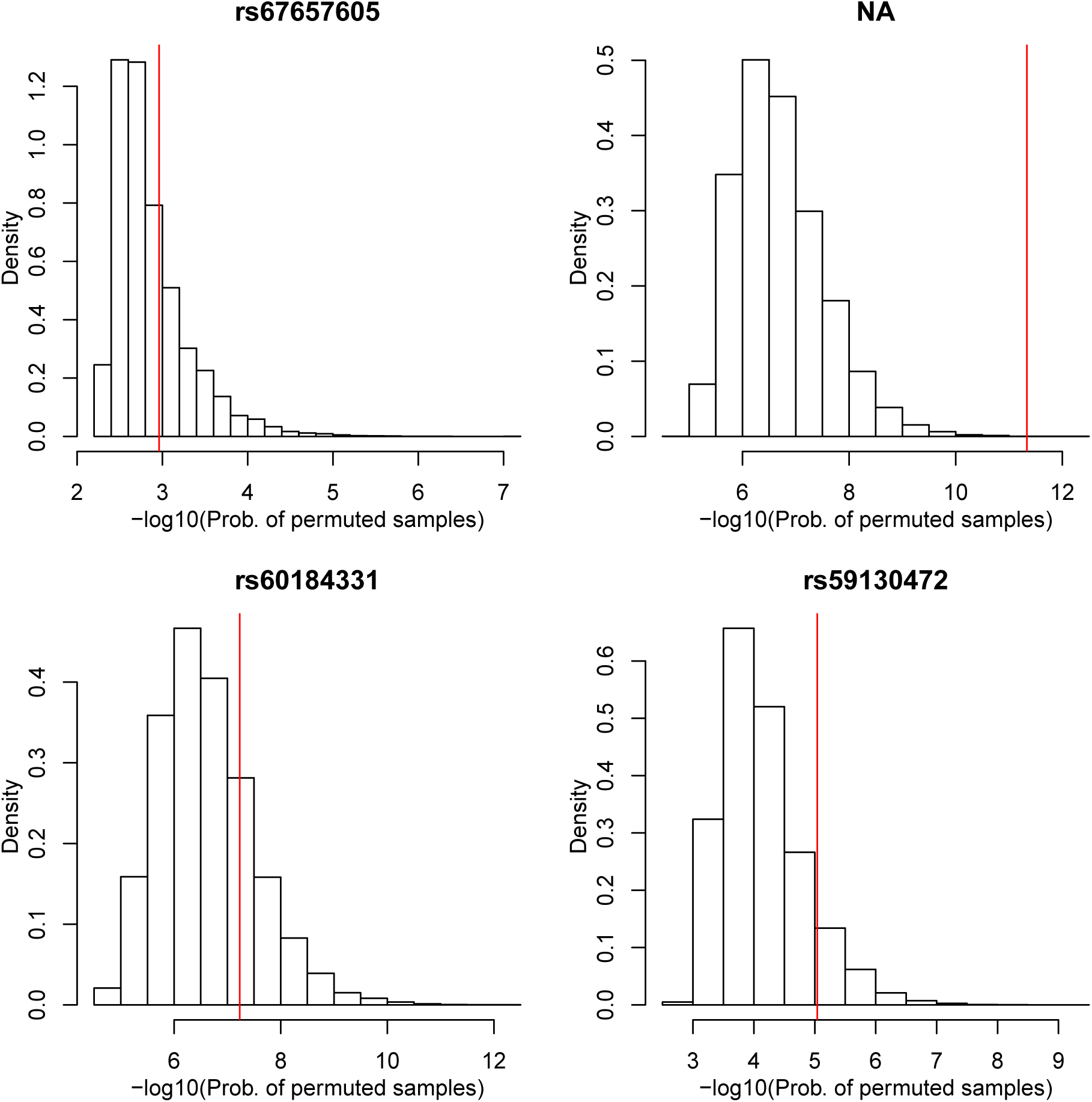
Distribution of the probability of permuted samples in an omnibus exact test for HWP and EAF with multi-allelic variants on the X chromosome.

### 4.2 Autosomal variants

We extracted multi-allelic variants of chromosome 7 of the JPT sample of the 1000 genomes project in order to illustrate the autosomal tests developed in Section 2.3. We used chromosome 7 because its size is similar to that of the X chromosome considered in the previous section. Multi-allelic variants on chromosome 7 are also rare. Of all 4.7M variants, 84.81% were monomorphic, 15.12% were bi-allelic, 0.06% were tri-allelic, and 0.004% had four or more alleles. Chromosome 7 has a larger percentage of bi-allelic variants than chromosome X. We consider a few examples of tri-allelic variants. Table 9 shows genotype counts and p-values of exact tests for six tri-allelic variants. These variants are registered as indels or SNVs that have at least two alternate alleles. The p-value of the omnibus exact test was in all cases estimated by a permutation procedure with 20.000 draws, in order to avoid the computational burden of the complete enumeration algorithm for autosomal variants. This gives an estimate of the p-value that is within 0.01 units of its true value with 99% confidence (Guo and Thompson, 1992). Standard HWP exact tests for all individuals and for males and females separately were carried out with a complete enumeration algorithm in the tri-allelic case (Louis and Dempster, 1987), and with a network algorithm for variants with more than three alleles (Engels, 2009).

**Table 9:**
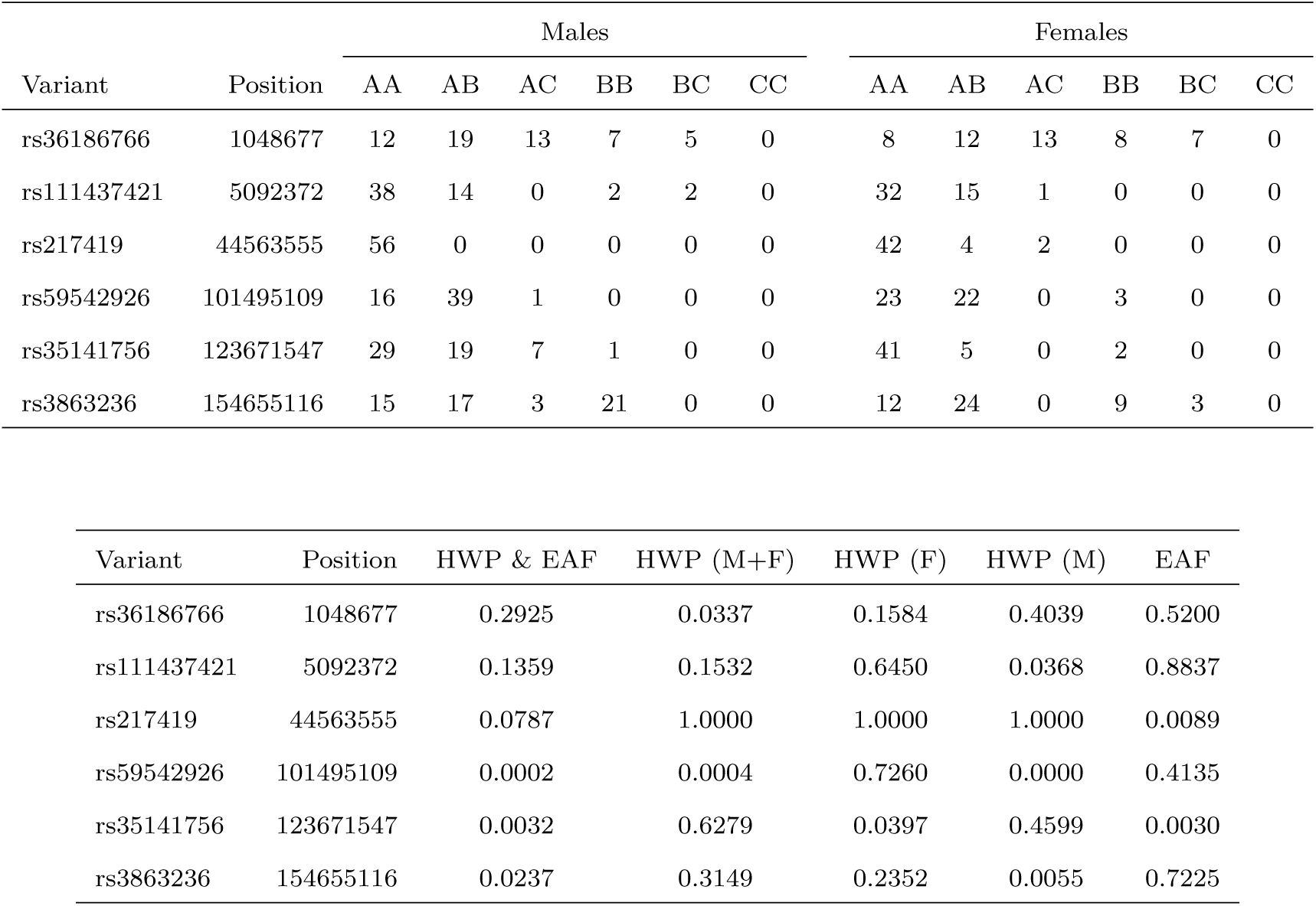
Variant, position (in bp), genotype counts and exact p-values for six tri-allelic variants on chromosome 7. HWP & EAF: omnibus exact test for HWP and EAF jointly, HWP (M+F): standard exact test with all individuals, HWP (F): standard exact test for HWP in females, HWP (M) standard exact test for HWP in males, EAF: Fisher exact test for equality of allele frequencies in males and females.

Variant rs36186766 is not significant in the joint HWP & EAF test, as assessed by its permutation p-value. This variant is significant in a standard exact test for HWP, but not when males and females are tested separately. This variant has an excess of heterozygotes, both in males and females, and this becomes significant when their genotype counts are summed. This apparently goes unnoticed in the omnibus test, as the latter considers the allele frequencies too.

Variant rs111437421 is significant only in a HWP test for males only. This goes unnoticed if the standard exact test on all individuals is used, and it also goes unnoticed in the proposed omnibus test, though the latter has a smaller p-value than the standard test.

Variant rs217419 is monomorphic in males, and has low frequencies of the alternate alleles in females. A standard exact test shows neither evidence for HWD overall, nor in males or in females. However, the allele frequencies in males and females do differ significantly, which is also reflected by a close to significant p-value in the omnibus test.

Variant rs59542926 is highly significant in both the omnibus and the standard HWP test. The deviation can be ascribed to males, which have an excess of heterozygotes.

Variant rs35141756 is clearly non-significant in a standard exact test for HWP that does not consider gender. There are however, significant differences in allele frequencies between males and females. Moreover, there is evidence that females deviate from HWP. The joint test (HWP & EAF) is significant in this case. This variant has a peculiar disequilibrium pattern and shows *attenuation of evidence* against HWP in the standard exact test. The variant has an excess of heterozygotes in males, but a lack of heterozygotes in females, which average out over all if the sexes are not distinguished.

Variant rs3863236 is significant in the omnibus test for HWP & EAF, but non-significant in a standard test for HWP. There is no evidence for differences of allele frequencies in the sexes, but males deviate significantly from HWP. In this case the joint tests reveals this, as it assumes HWP in both sexes. This variant also shows signs of attenuation of evidence for the standard exact test for HWP. The standard p-value of a test for HWP is larger than the p-values of the separate tests in males and females, despite the fact that this test has more power due to a doubled sample size. In this case for males there are fewer heterozygotes than expected, whereas for females there are more heterozygotes than expected.

We continue to study all 2992 tri-allelic variants on chromosome 7 simultaneously. Figure 3 shows some graphics summarizing these variants. Figure 3A shows that tri-allelics do occur all along chromosome 7, and that disequilibrium mostly arises from an excess of heterozygotes. We note that due to the use of the permutation test, the p-values are bounded above by −*log*_10_(1/20.000) = 4.3. For variants that had no permuted samples with smaller probabilities than the observed sample, the permutation p-value was set to this limit. Some of the tri-allelic variants apparently do have a p-value smaller than the Bonferroni limit, but this would become visible only at a greatly increased computational cost. The limit on the number of permutations also explains the truncation observed in the QQ-plot at 4.3 in Figure 3B. Like the X chromosomal tri-allelics previously studied in Section 4.1, the observed heterozygosity is often larger than the expected heterozygosity, with 68% of the variants are above the line y = x, showing that the tri-allelics on chromosome 7 are also characterized by a general excess of heterozygosity. Figure 3C also shows that variants with extreme observed heterozygosities often have significant differences in allele frequencies between the sexes.

**Figure 3:**
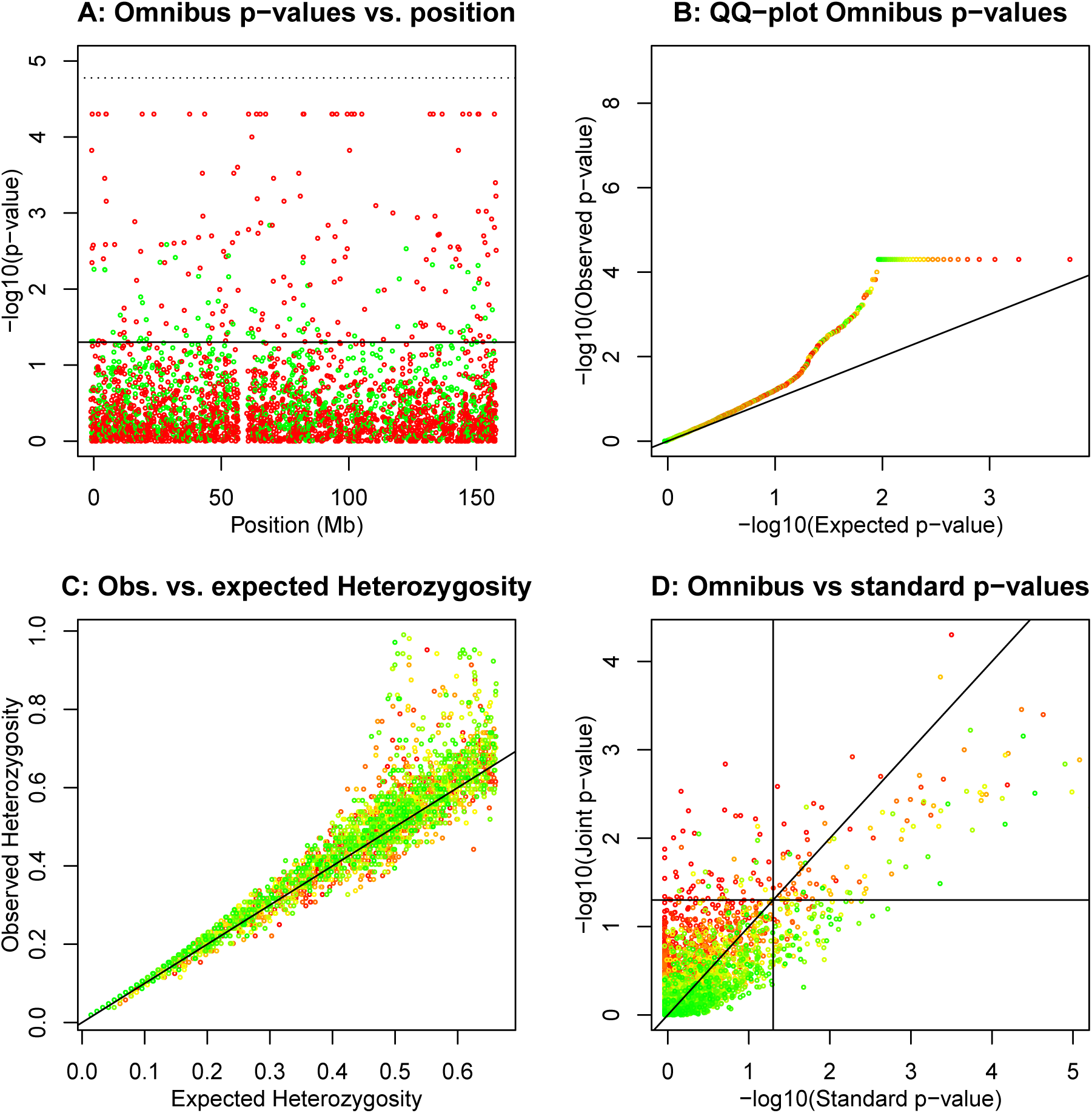
Hardy-Weinberg equilibrium tests for 2992 tri-allelic variants on chromosome 7. A: p-values of the omnibus exact test versus variant position. The solid horizontal line corresponds to *α* = 0.05, dotted horizontal line corresponds to the Bonferroni corrected threshold. Variants colour-coded according to having excess of heterozygotes (red) or lack of heterozygotes (green) with respect to HWP. B: QQ-plot of the omnibus exact p-values. C: Observed versus expected heterozygosity. D: Omnibus exact p-values versus exact p-values obtained in a test using the females only. Panels B, C and D have variants colour-coded according to the p-value of an exact test for EAF (significant to non-significant from red to green). Horizontal and vertical lines correspond to *α* = 0.05.

We finish the analysis of chromosome 7 with a few examples of variants that have more than three alleles. Genotype counts of four, five and six allelic variants are given, stratified by sex, in Table 10.

**Table 10:**
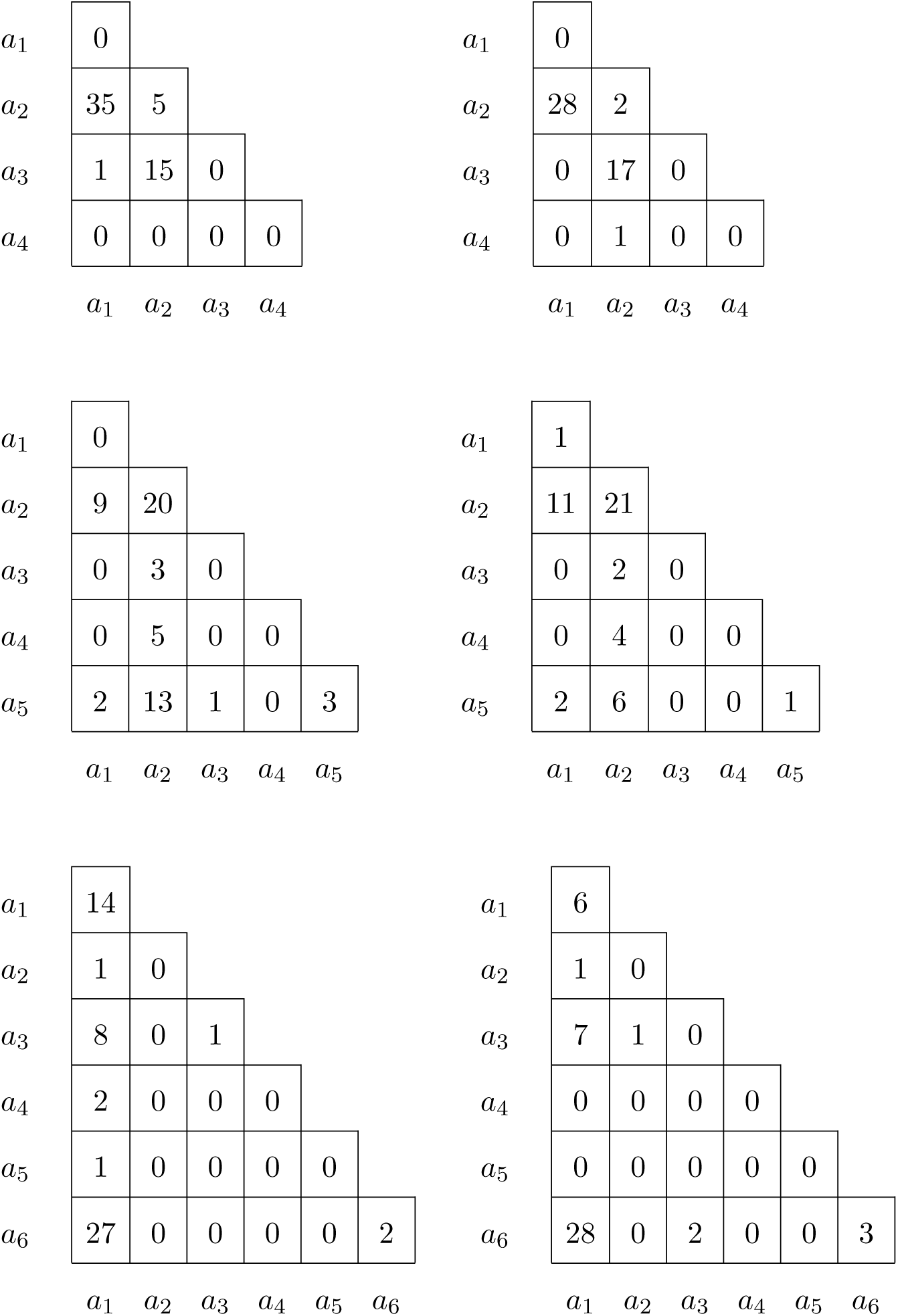
Genotype counts for males (left) and females (right) of a four, five and six allelic variants on chromosome 7.

|  |  |  |  |  |
| --- | --- | --- | --- | --- |
| $a_1$ | 0 | | | |
| $a_2$ | 35 | 5 | | |
| $a_3$ | 1 | 15 | 0 | |
| $a_4$ | 0 | 0 | 0 | 0 |
| | $a_1$ | $a_2$ | $a_3$ | $a_4$ |
| $a_1$ | 0 | | | |
| $a_2$ | 28 | 2 | | |
| $a_3$ | 0 | 17 | 0 | |
| $a_4$ | 0 | 1 | 0 | 0 |
| | $a_1$ | $a_2$ | $a_3$ | $a_4$ |

|  |  |  |  |  |  |
| --- | --- | --- | --- | --- | --- |
| $a_1$ | 0 | | | | |
| $a_2$ | 9 | 20 | | | |
| $a_3$ | 0 | 3 | 0 | | |
| $a_4$ | 0 | 5 | 0 | 0 | |
| $a_5$ | 2 | 13 | 1 | 0 | 3 |
| | $a_1$ | $a_2$ | $a_3$ | $a_4$ | $a_5$ |
| $a_1$ | 1 | | | | |
| $a_2$ | 11 | 21 | | | |
| $a_3$ | 0 | 2 | 0 | | |
| $a_4$ | 0 | 4 | 0 | 0 | |
| $a_5$ | 2 | 6 | 0 | 0 | 1 |
| | $a_1$ | $a_2$ | $a_3$ | $a_4$ | $a_5$ |

|  |  |  |  |  |  |  |
| --- | --- | --- | --- | --- | --- | --- |
| $a_1$ | 14 | | | | | |
| $a_2$ | 1 | 0 | | | | |
| $a_3$ | 8 | 0 | 1 | | | |
| $a_4$ | 2 | 0 | 0 | 0 | | |
| $a_5$ | 1 | 0 | 0 | 0 | 0 | |
| $a_6$ | 27 | 0 | 0 | 0 | 0 | 2 |
| | $a_1$ | $a_2$ | $a_3$ | $a_4$ | $a_5$ | $a_6$ |
| $a_1$ | 6 | | | | | |
| $a_2$ | 1 | 0 | | | | |
| $a_3$ | 7 | 1 | 0 | | | |
| $a_4$ | 0 | 0 | 0 | 0 | | |
| $a_5$ | 0 | 0 | 0 | 0 | 0 | |
| $a_6$ | 28 | 0 | 2 | 0 | 0 | 3 |
| | $a_1$ | $a_2$ | $a_3$ | $a_4$ | $a_5$ | $a_6$ |

The different test results for these variants are given in Table 11. The first variant in this table (without RS identifier) has four allies. Almost all alleles occur in heterozygotes. Homozygotes for the reference allele (*a*_1_) are missing. There is a very strong heterozygote excess and consequently all exact tests except the one for EAF are highly significant. The results suggest this variant has genotyping problems.

**Table 11:** Test results for three multi-allelic variants on chromosome 7 of the JPT sample.

|  | Variant | # alleles | HWP & EAF | HWP (M+F) | HWP (M) | HWP (F) | EAF |
| --- | --- | --- | --- | --- | --- | --- | --- |
| 1 | NA | 4 | 0.0000 | 0.0000 | 0.0000 | 0.0000 | 0.6964 |
| 2 | rs145685769 | 5 | 0.9483 | 0.7216 | 0.9285 | 0.9505 | 0.2968 |
| 3 | rs71774128 | 6 | 0.0035 | 0.0005 | 0.0571 | 0.0044 | 0.3520 |

Variant rs145685769 is consistently non-significant in all exact tests applied, and therefore seems compatible with HW proportions and equality of allele frequencies in the sexes.

Variant rs71774128 is significant in all exact HWP tests. This variant has a heterozygosity that is larger than expected under HWP, both for males and females.

## 5 Discussion

In this paper we have developed exact test procedures for HWP that take gender into account, for variants with multiple alleles. We have illustrated these procedures with indels, though are procedures are equally relevant for micro-satellites which are widely used in molecular ecology.

In the case of X chromosomal variants it seems compelling to use sex in the analysis. If, as has been common practice until recently, only the data from the females are used, then the number of X chromosomes in the sample will decrease by one third. Consequently, estimates of allele frequencies will be less precise, to the detriment of all the statistical analyses that follow. Moreover, as we have pointed out previously (Graffelman and Weir, 2016), null alleles can go undetected in heterozygote form in females, but show up in males. For the X chromosome, a test that accounts for sex, and tests HWP and EAF jointly, therefore seems an attractive option.

For the autosomes, taking gender into account seems a priori, not necessary, and the standard practice is to use the total genotype counts in tests for HWP. There is no loss of data by not considering sex in this case. However, as the examples in Section 4.2 show, unexpected genotype count patterns are sometimes detected if gender is considered. Such patterns may represent chance effects, or may be the result of some genotyping problem. It therefore seems sensible to at least use tests for equality of allele frequencies in the sexes as part of the quality control process. We do not suggest replacing the standard autosomal exact test for HWP, widely used in quality control, by the proposed exact procedures that take sex into account. However, we do suggest that significant autosomal GWAS findings could be checked for unexpected patterns by the autosomal procedures proposed in this paper.

For bi-allelic variants, it is well-known that the classical chi-square test is problematic at low minor allele frequencies, due to low expected counts for the minor homozygote. For variants with multiple alleles the asymptotic chi-square test is even more problematic, because typically some of the alleles have low frequencies. Taking gender into account in tests for HWP with multiple alleles further aggravates the sparseness problem, making it even more difficult to apply chi-square tests that rely on asymptotic results. Given a fixed total sample size, if one wishes to account for gender this implies *k* extra categories in the X-chromosomal case, whereas it doubles the number categories for the autosomal case, and inevitably more categories with lower counts appear. Exact procedures are therefore indicated in this setting.

Efficient algorithms for calculating HW exact test probabilities for the bi-allelic case have been proposed by Wigginton et al. (2005) and Chang et al. (2015). The bi-allelic exact procedure for testing HWP and EAF jointly at X-chromosomal variants is computationally feasible for complete X chromosomes with Chang’s algorithm implemented in both PLINK 2.0 and R-package HardyWeinberg (Graffelman, 2015). Accounting for gender for variants with multiple alleles augments the number of possible outcomes for the exact test, and implies an increase of the computational burden with respect to the standard procedures that do not take gender into consideration. All computational improvements that have been proposed for multiple alleles such as Huber’s (2006) faster generation of permutations, the Markov-chain approach from Guo and Thompson (1992), and the network algorithm of Engels (2009) could be used to reduce the amount of computation involved, but are left unexplored here. Most of the variants of the 1000 Genomes project are bi-allelic, and for tri-allelics, the enumeration algorithm was feasible for the studied sample size. The computational burden depends on the distribution of the allele counts. Often, alleles beyond the most frequent alternate allele are rare, and for these cases not much extra computation is required. In the case of uniformly distributed allele counts (with close to maximal expected heterozygosity) the number of possible outcomes is much larger, and this increases the computational cost of the enumeration algorithm.

In the exact test with multiple alleles, tied outcomes (with the same probability) can easily arise (see the example in Table 4). Tied outcomes can provoke inexact p-values, as on the computer such tied outcomes might have a slightly different probability due to rounding errors. Let *p*(*o*) represent the probability of the observed sample. The rounding problem with ties can be avoided if outcomes are considered to have a smaller probability than the observed sample whenever their probability is smaller than *p*(*o*) + *ε*, with *ε* some convenient small number. Differences in exact p-values across computer programs are likely to be due to ties.

The analysis of tri-allelic variants of the JPT data shows that deviation from HWP is often due to an excess of heterozygotes, both for the X chromosome and for chromosome 7. This has also been described for the far more common bi-allelic variants (Graffelman <u>et al.</u>, 2017), and was postulated to be a consequence of polymorphism duplication. The analysis of the tri-allelics also reveals that HWD often goes together with a difference in allele frequencies between the sexes.

Some of the autosomal examples in the paper show that in a standard exact test for HWP, *attenuation of evidence for HWD* can occur if one sex has a deficiency of heterozygotes, whereas the other sex has an excess. Such opposing heterozygosities can average out, such that disequilibrium goes unnoticed when sex is not considered. The proposed omnibus test seems able to detect these cases.

## Acknowledgments

This work was partially supported by grant 2014SGR551 from the Agència de Gestió ďAjuts Universitaris i de Recerca (AGAUR) of the Generalitat de Catalunya, by grant MTM2015-65016-C2-2-R (MINECO/FEDER) of the Spanish Ministry of Economy and Competitiveness and by and by grant R01 GM075091 from the United States National Institutes of Health.

The authors declare that there are no conflicts of interest.

